# Structural basis of substrate recognition and transport in bacterial ACS transporters

**DOI:** 10.64898/2026.05.20.726482

**Authors:** Kim Bartels, Katharina E. J. Jungnickel, Josi Steinke, Shindhuja Joel, Christian Löw

**Affiliations:** Centre for Structural Systems Biology (CSSB), Notkestraße 85, 22607 Hamburg, Germany; European Molecular Biology Laboratory (EMBL) Hamburg, Notkestraße 85, 22607 Hamburg, Germany; University Medical Centre Hamburg-Eppendorf (UKE), Hamburg, Germany

**Keywords:** MFS, SLC17, symporter, LgoT, DgoT, GarP, GudP, ExuT, proton-coupling, LCP, X-ray structure, ACS transporter

## Abstract

Membrane transporters of the major facilitator superfamily (MFS) mediate uptake of diverse metabolites, yet the molecular basis of substrate recognition within many bacterial families remains unclear. The anion:cation symporter (ACS) family is conserved from bacteria to humans and includes medically relevant solute carrier transporters, but only few bacterial members have been functionally characterized. Here, we combine genetics, biochemistry, transport assays, and structural biology to define the substrate specificity of five ACS transporters from *Escherichia coli*. Using systematic growth complementation assays in deletion strains, we assign physiological substrates to each transporter, identifying DgoT, LgoT, and ExuT as specific uptake systems for D-galactonate, L-galactonate, and galacturonate/glucuronate, respectively, while revealing overlapping roles for GarP and GudP in C6 sugar acid uptake. NanoDSF ligand binding assays show highly selective recognition of sugar acids but do not predict transport activity. Proton-coupled uptake was directly demonstrated using reconstituted proteoliposomes, defining strict stereoselectivity of LgoT and DgoT. We determined the 2.2 Å X-ray structure of LgoT in an inward-open conformation, revealing a canonical MFS fold with a conserved but differentially tuned substrate-binding cavity. Comparative structural and sequence analyses across ACS members identify a conserved core for coordination of sugar acid carboxylate and hydroxyl groups, while localized substitutions modulate steric and electrostatic properties to enable discrimination of substrate size and stereochemistry. These results provide a framework for substrate recognition in bacterial ACS transporters and establish LgoT as a structural model for stereo-selective proton-coupled organic anion transport.

## Introduction

Membrane transporters of the major facilitator superfamily (MFS) mediate the selective uptake and export of small molecules across biological membranes, shaping nutrient utilization, cellular signaling, and metabolic homeostasis [1-3]. One clade within this superfamily, the anion:cation symporter (ACS) family, is highly conserved from bacteria to humans and is best known for transporting organic anions in symport with protons or sodium ions [4, 5]. In mammals, ACS proteins constitute the solute carrier family 17 (SLC17), whose members participate in processes such as renal organic anion secretion, lysosomal catabolism of glycoconjugates, and synaptic neurotransmitter loading [5-9].

SLC17 transporters are divided into four groups [5, 10]: (i) type I phosphate transporters (NPT1, NPT3, NPT4, NPT5), originally thought to transport phosphate in a sodium dependent manner, but now recognized primarily as organic anion carriers involved in urate metabolism and drug secretion; (ii) sialin, a lysosomal exporter of sialic acid, glucuronic acid, and related acidic sugars; (iii) vesicular glutamate transporters VGLUT1–3, which specifically load glutamate into synaptic vesicles; and (iv) the vesicular nucleotide transporter VNUT, responsible for ATP loading into secretory vesicles. Defects in members of this family are implicated in various diseases such as hyperuricemia, gout, and lysosomal storage disorders [11-18].

The bacterial ACS transporter DgoT, which mediates D-galactonate uptake, has emerged as a powerful model for investigating the atomistic basis of proton-coupled transport in the SLC17 family due to its high sequence and functional similarity to mammalian homologs [19-22]. Structural studies of DgoT, followed by rat VGLUT2 and human sialin, revealed the canonical 12-transmembrane (TM) MFS fold, consisting of N-terminal (TM1–6) and C-terminal (TM7–12) bundles connected by a cytoplasmic intracellular helical (ICH) domain, forming a central substrate-binding cavity [21, 23-26]. Combined structural and molecular dynamics analyses elucidated how DgoT coordinates the carboxylate and hydroxyl groups of D-galactonate and couples substrate binding to proton translocation [19]. Comparisons with VGLUT2 and sialin revealed conserved principles of carboxylate recognition and family-specific adaptations, including differences in proton-binding sites that modulate proton-to-substrate stoichiometry [8, 24-26]. In contrast, NPT transporters lack these proton-binding residues, consistent with their sodium-dependent transport mechanism.

Many bacterial ACS proteins are predicted to transport anionic sugars or phthalates based on their operon structure [27-34], but only a subset has been biochemically characterized. *Escherichia coli* (*E. coli*) encodes six ACS transporters (ExuT, RhmT, DgoT, LgoT, GarP and GudP), all proposed to function as proton-coupled anionic sugar symporters [5] (Figure 1A). This repertoire enables *E. coli* to utilize diverse sugar acids, including D-galactonate, L-galactonate, D-gluconate, D-glucuronate, D-galacturonate, and D-glucarate, as sole carbon and energy sources [27, 35-37]. These sugar acids are abundant in the mammalian gut as products of host metabolism, dietary polysaccharide degradation, and microbial oxidation reactions [38-41]. Catabolism funnels them into central metabolism via intermediates such as pyruvate [27-29, 36, 37]. In many cases, the substrate itself acts as the operon inducer [27, 28, 31, 32, 39].

**Figure 1:**
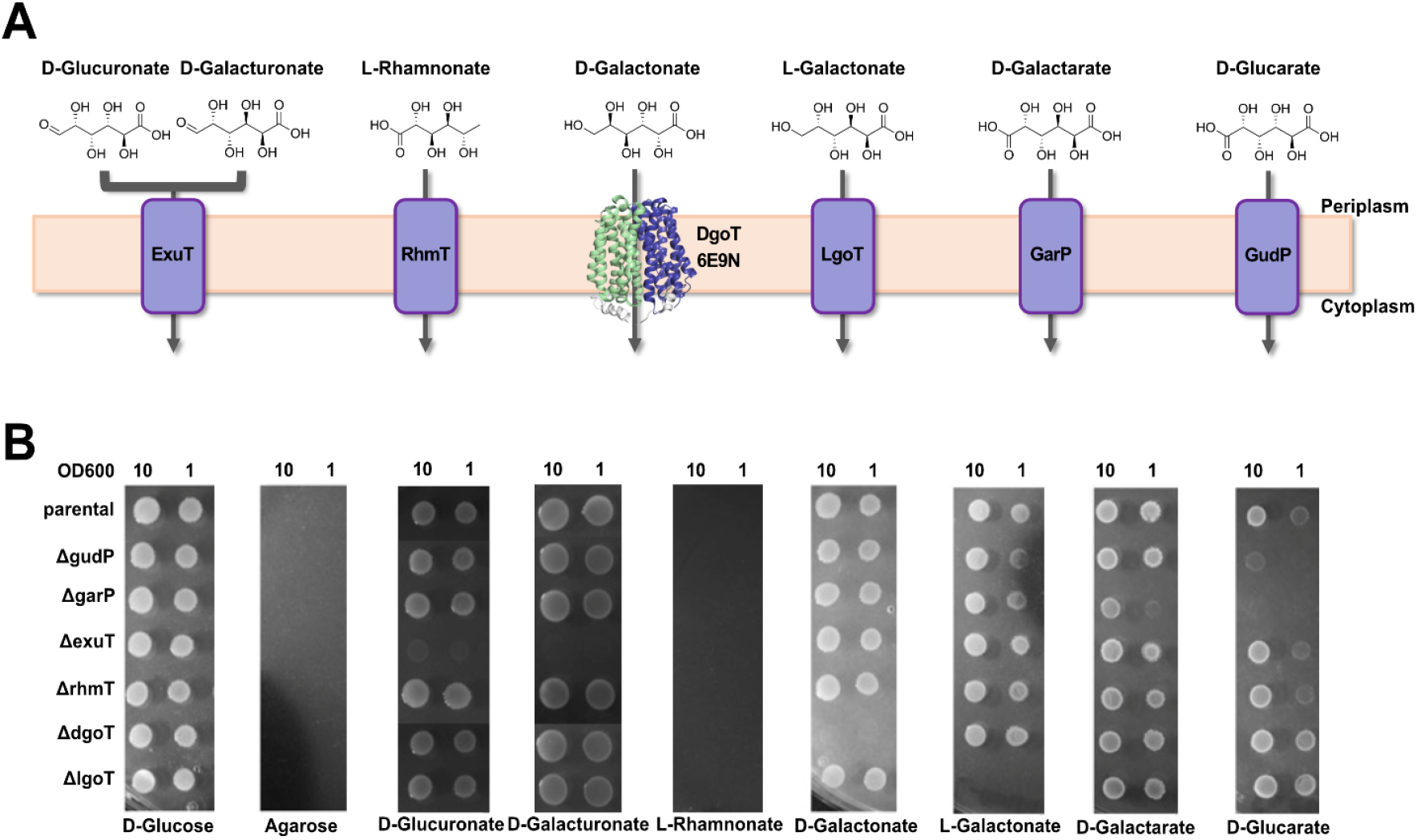
*In vivo* growth assay. (A) *E. coli* ACS transporters with suggested substrates are illustrated. An experimentally determined structure is available for DgoT (PDB-ID 6E9N) [21]. (B) Growth of *E. coli* on agar plates supplied with different sugars as carbon sources. Different *E. coli* parental and knockout strains were cultured in minimal medium, washed and spotted onto agar plates at an OD_600nm_ of 10 and 1. Growth of colonies was assessed after 24 h incubation at 37°C.

DgoT remains the best-characterized bacterial ACS transporter. It selectively transports D-galactonate [21], and structural studies in inward- and outward-open states revealed the molecular basis of substrate recognition and proton coupling [20, 21]. In contrast, far less is known about LgoT, the proposed transporter for L-galactonate. The lgo operon contains three genes: lgoT (yjjL), yjjM, and yjjN. These genes are all strongly upregulated when *E. coli* grows on L-galactonate as the sole carbon source [31]. YjjM functions as a pathway regulator, while YjjN is an oxidoreductase converting L-galactonate to D-tagaturonate, the first dedicated catabolic step [31, 37]. Unlike D-galactonate, whose metabolism follows a dedicated pathway, L-galactonate catabolism merges with the D-galacturonate pathway [34, 42] (SI Figure 1). Despite functional predictions, direct biochemical validation and structural insights into LgoT have been lacking.

Here, we present a comprehensive functional analysis of the members of the *E. coli* ACS transporters, with a structural focus on LgoT. Through a combination of ligand binding, substrate transport and *in vivo* growth assays we assigned substrates to each member, confirming LgoT as the L-galactonate transporter and revealing the broad sugar acid specificity of the ACS family. We also report the high-resolution X-ray structure of LgoT, highlighting conserved architectural features and unique molecular determinants underlying selective recognition. Comparative analysis with DgoT shows how subtle changes in the binding pocket enable discrimination between epimeric sugar acids. Together, our findings expand the mechanistic framework of ACS transport, define the molecular basis of substrate specialization, and illustrate how *E. coli* leverages ACS diversity to achieve metabolic flexibility. More broadly, LgoT complements DgoT as a model system for studying proton-coupled organic anion transport and provides a structural foundation for future investigations of ACS function across bacteria and eukaryotes.

## Results and Discussion

### Substrate assignment of the *E. coli* ACS transporter family by *in vivo* growth assays

To define the physiological substrates of the six *E. coli* ACS transporters (ExuT, RhmT, DgoT, LgoT, GarP and GudP), we performed growth complementation assays using KEIO single-gene deletion strains, which are derived from the *E. coli* K-12 background [43], on a panel of 24 different carbon sources. The parental strain BW25113 and the corresponding Δ*exuT*, Δ*rhmT*, Δ*dgoT*, Δ*lgoT*, Δ*garP* and Δ*gudP* mutants were first cultivated in minimal medium supplemented with glucose. After glucose removal and normalization to equal cell density, dilution series of each strain were spotted onto minimal agar plates in which a single sugar or sugar acid served as the sole carbon source (Figure 1B), and growth was assessed.

First, we established that none of the used strains was able to sustain growth on the polysaccharide agarose alone, which was present in all used agar plates. As expected from the metabolic defect characteristic of the K-12 strain [44-47], neither the parental nor any knockout strains grew on D-lactose, L-arabinose, or L-rhamnose (Table 1). Similarly, none of the strains utilized D-sucrose, D-raffinose, meso-xylitol, or myo-inositol, consistent with known limitations of K-12 derivatives. In contrast, robust growth was observed for all strains on D-fructose, D-glucose, D-galactose, D-maltose, the sugar acid D-gluconate, D-ribose, N-Acetyl-D-glucosamine, D-sorbitol, and D-trehalose indicating that none of the tested ACS transporters are required for uptake of these substrates (Table 1).

**Table 1:**
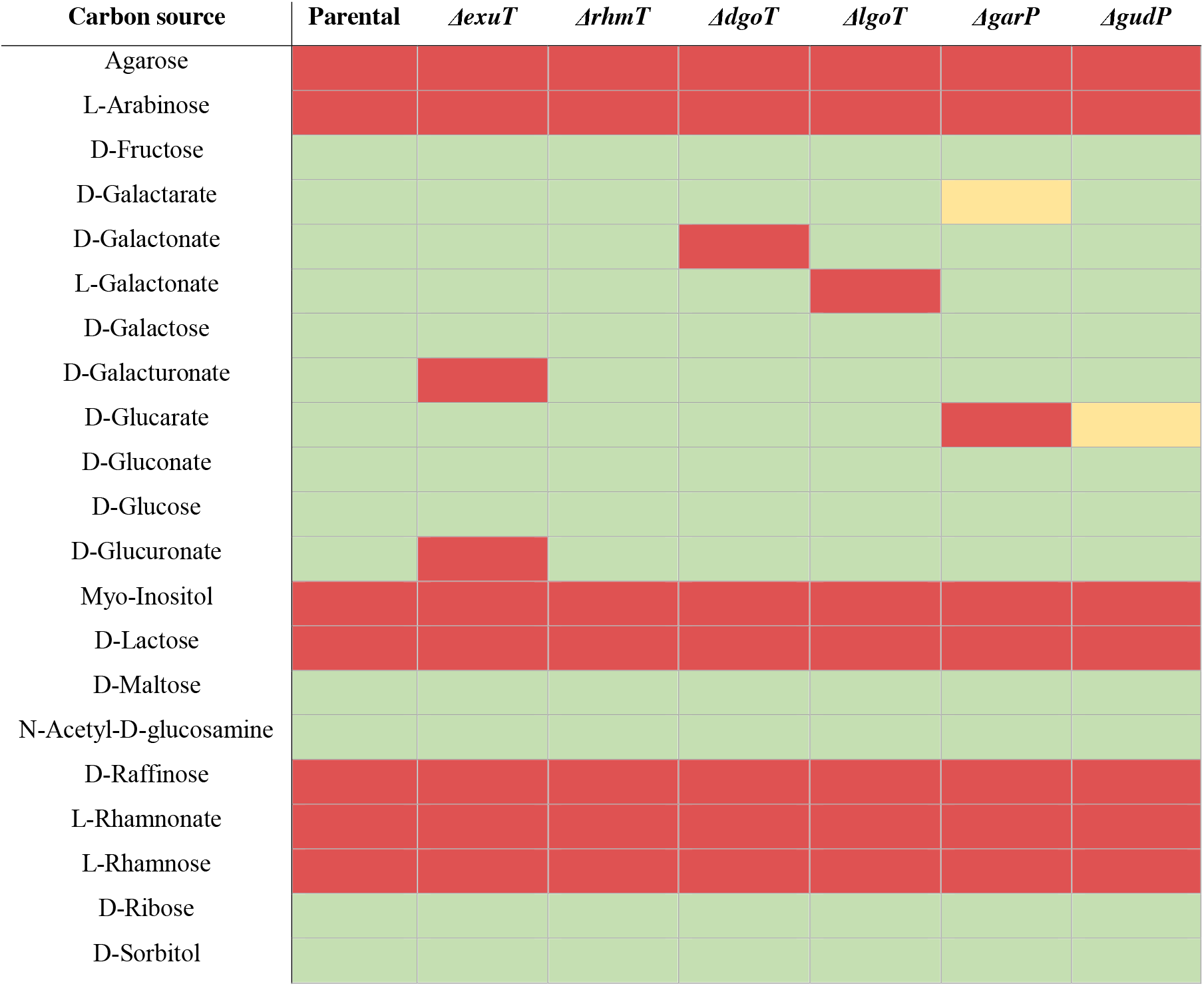
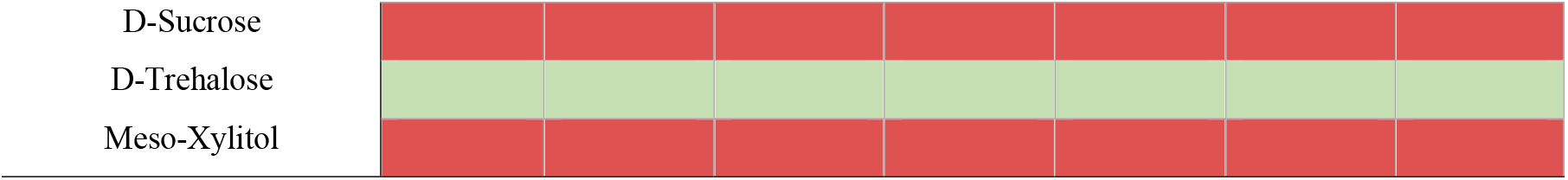
Summary of the ability of K-12 *E. coli* parental and ACS transporter knockout strains to grow on different carbon sources. Red denotes no growth, green denotes growth, yellow denotes impaired growth.

Distinct transporter dependent phenotypes emerged on the six sugar acids D-galactonate, L-galactonate, D-galactarate, D-glucarate, D-galacturonate, and D-glucuronate (Figure 1, Table 1). Δ*dgoT* exhibited a complete loss of growth on D-galactonate, while Δ*lgo*T failed to grow on L-galactonate. Δ*exuT* showed abolished growth on both D-galacturonate and D-glucuronate. These knockout-specific phenotypes directly assign DgoT, LgoT, and ExuT as the sole transporters required for D-galactonate, L-galactonate, and galacturonate/glucuronate utilization, respectively, consistent with previous predictions or biochemical observations.

More complex patterns were observed for the remaining two ACS members, GarP and GudP, both implicated in the uptake of C6 dicarboxylate sugar acids. Based on the operon structure and earlier reports, GarP has been proposed to transport both D-galactarate and D-glucarate [29, 30]. In our assay, Δ*garP* failed to grow on D-glucarate and showed markedly reduced growth on D-galactarate (Figure 1, Table 1). Conversely, Δ*gudP*, predicted to be the primary D-glucarate transporter [29, 30], displayed measurable growth impairment on D-glucarate but normal growth on D-galactarate. These data reveal overlapping yet asymmetric substrate profiles for these transporters. Importantly, the D-galactarate and D-glucarate metabolism converge into a shared catabolic pathway, differing only at substrate-specific dehydrogenase steps [29], and both the *garP* and *gudP* operons are induced by D-galactarate and D-glucarate, with stronger induction levels by D-galactarate [32]. These regulatory and metabolic features provide an explanation for the observed phenotypes. Based on these findings, we propose that GarP is the primary transporter for both D-galactarate and D-glucarate, whereas GudP provides secondary, partially redundant transport capacity for D-glucarate. In the absence of *gudP*, cells grow on both substrates through uptake *via* GarP but with measurable growth impairment on D-glucarate, indicating that both transporters contribute to D-glucarate uptake under physiological conditions. In the absence of *garP*, induction of a different transport system can partially rescue D-galactarate uptake but is insufficient to sustain normal growth. If this alternate transport system is GudP cannot be concluded from the current data.

The substrate of RhmT could not be established. None of the strains, including the parental strain BW25113, grew on L-rhamnonate, the anticipated substrate (Figure 1, Table 1). Given that K-12 derivatives lack the enzymes required for L-rhamnose catabolism, it is likely that steps necessary for L-rhamnonate metabolism are also absent or not expressed, precluding substrate assignment for RhmT in this context. Interestingly, while deletion of ACS transporters affected growth on D-galactonate, the aldonic acid of D-galactose, growth was unaffected on D-gluconate, the aldonic acid of D-glucose (Figure 1, Table 1). This is consistent with the presence of several non-ACS MFS transporters dedicated to D-gluconate uptake [48]. In summary, growth phenotypes on defined carbon sources enabled initial substrate assignment of the ACS family members except for RhmT.

### *In vitro* binding assays of ACS transporters by nanoDSF

To biochemically characterize the *E. coli* ACS transporter family, each of the six members was cloned with an affinity tag and recombinantly expressed in *E. coli* (SI Figure 2A). All transporters were expressed at reasonable yields; however, only DgoT, LgoT, GarP, and GudP remained homogeneous and stable following detergent solubilization and purification (SI Figure 2B). ExuT and RhmT showed low stability and pronounced aggregation and were excluded from further *in vitro* analysis.

Guided by the substrate assignments obtained from the *in vivo* growth assays, we assessed ligand binding using nano-differential scanning fluorimetry (nanoDSF). Binding-induced stabilization was monitored as an increase in the melting temperature (T_m_) of the transporter–ligand complex (Figure 2A). To broadly interrogate ligand specificity, we screened 37 compounds at a concentration of 2.5 mM, including sugars, sugar acids, sugar alcohols, amino sugars, and selected amino acids (SI Figure 3). The latter were included because several mammalian SLC17 homologs, such as VGLUTs and sialin, are known to also transport acidic amino acids.

**Figure 2:**
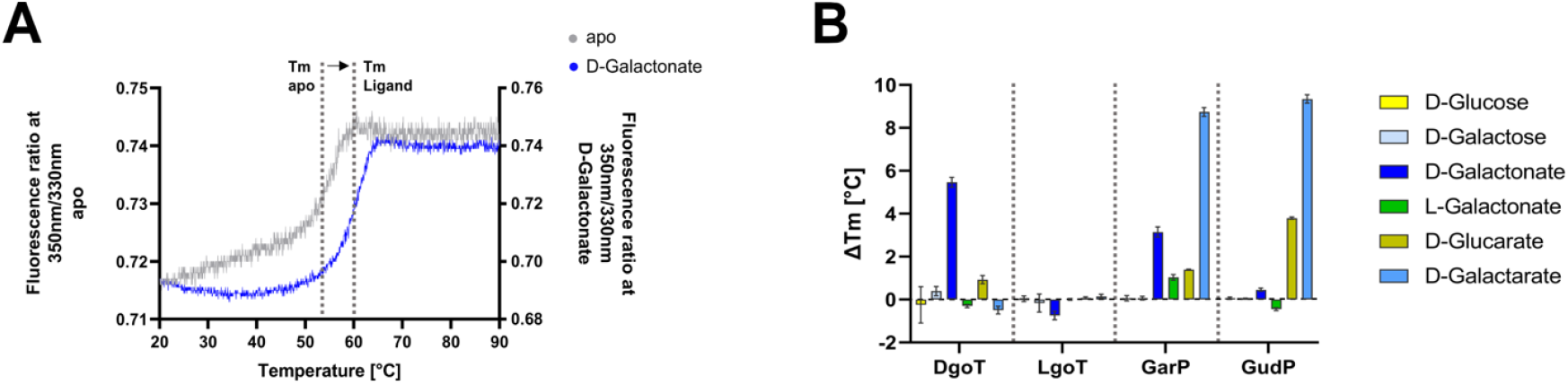
Ligand binding of ACS transporter from *E. coli*. (A) Thermal unfolding curve of DgoT without ligand addition (apo) or with ligand (D-galactonate) as example of a thermal stability measurement using nanoDSF. The samples are heated from 20 °C to 90 °C while the fluorescence at 330 nm and 350 nm for tyrosine and tryptophan residues respectively is recorded. Plotting the ratio of the fluorescence at 350 nm and 330 nm against the temperature results in an unfolding curve, with its inflection point defined as the melting temperature (T_m_). If ligand binding stabilizes the protein, this leads to an increase in T_m_. (B) Difference of T_m_ upon ligand addition to DgoT, LgoT, GarP and GudP. Stabilities of the proteins were measured at a concentration of 0.5 mg/mL with ligands added at a concentration of 2.5 mM. Selected sugars are color-coded. Error bars correspond to the standard deviation (n = 3).

Each of the four screened transporters exhibited a remarkable narrow ligand specificity. GarP was stabilized by D-galactarate (ΔT_m_ = 8.7 ± 0.2°C), D-galactonate (ΔT_m_ = 3.1 ± 0.2°C), and to a lesser extent by D-glucarate (ΔT_m_ = 1.4 ± 0.2°C). GudP showed pronounced stabilization with D-galactarate (ΔT_m_ = 9.3 ± 0.2°C) and D-glucarate (ΔT_m_ = 3.8 ± 0.1°C). DgoT was selectively stabilized by D-galactonate (ΔT_m_ = 5.5 ± 0.2°C) only (Figure 2B). In contrast, LgoT did not show detectable stabilization in the presence of any tested compound, including its predicted substrate L-galactonate (Figure 2B).

To verify that the observed thermal shifts reflect specific ligand interaction, we performed concentration-dependent nanoDSF measurements to derive apparent dissociation constants (K_D,app_) [49, 50]. Dose-dependent stabilization was observed for DgoT with D-galactonate (K_D,app_ = 4.1 ± 1.2 mM); for GarP with D-galactarate (K_D,app_ = 5.0 ± 3.6 mM) and D-galactonate (K_D,app_ = 3.0 ± 0.7 mM); and for GudP with D-galactarate (K_D,app_ = 0.9 ± 0.4 mM) and D-glucarate (K_D,app_ = 3.8 ± 1.4 mM), confirming specific ligand engagement (SI Figure 4).

Because recombinant LgoT did not respond to L-galactonate, we tested whether pH or the membrane-mimetic environment affects ligand interaction. The rationale behind this approach is the fact, that substrate transport of LgoT is proton-coupled, implying that protonation of critical residues is crucial for substrate recognition and transport. Furthermore, a chosen membrane mimetic can have a strong influence on the conformation of a transporter. Mimetics, that rather favor the inward-open/release state of a transporter are not beneficial. Across the pH-range of 6.0–8.0, no stabilization was observed (SI Figure 5A-B). Purification of LgoT in alternative detergents (shorter chain detergents such as DM, NM, or a DDM/LDAO mixture), which decreased the apo T_m_ (44.7–51.1°C compared with 62.3°C in DDM), likewise revealed no ligand-induced stabilization (SI Figures 2C-E and 5C-D). Finally, LgoT was reconstituted into POPS-containing Salipros to restore a more native bilayer context (SI Figure 2F) [51-53]. This approach has been shown previously to expand the conformational landscape of a bacterial peptide transporter, allowing them to sample different conformations in this environment [54].

Even under these conditions, L-galactonate did not increase the melting temperature of LgoT, but the lipid environment had an overall strong influence on the thermostability of the transporter (SI Figure 5C). Taken together, these results highlight that while genetic evidence strongly supports LgoT as the L-galactonate transporter, ligand binding does not produce a measurable stabilizing effect under the tested *in vitro* conditions. This may reflect intrinsically weak thermal stabilization upon binding or conformational states in detergent or Salipros that are not thermally sensitive to ligand engagement. As a result, functional transport assays are required to reconcile this discrepancy.

Overall, nanoDSF binding analyses validated the specificity of DgoT, GarP, and GudP for their physiologically relevant sugar acids, but substrate binding does not automatically translate into transport.

### Liposome-Based Transport Assays Define Substrate Specificity of LgoT and DgoT

To distinguish substrate binding from transport, we assessed proton-coupled uptake of the ACS transporter LgoT and DgoT, reconstituted in proteoliposomes and loaded with the membrane-impermeable pH-sensitive dye pyranine. Transport was energized by generating an inwardly negative membrane potential. Liposomes composed of POPE:POPG (3:1, w/w), that mimic the inner membrane of *E. coli*, were loaded with 120 mM K^+^, exchanged into Na^+^ buffer, and then treated with the potassium-selective ionophore valinomycin to dissipate the K^+^ gradient. This created an inwardly negative membrane potential, which is then utilized by ACS transporters for proton-coupled substrate transport into the proteoliposomes. In turn, the influx of protons leads to a decrease of the proteoliposomes’ internal pH. Because pyranine fluorescence responds to intraluminal pH change, proton influx mediated by the symport of the ACS transporters could be monitored ratiometrically upon excitation at 415/460 nm (Figure 3A).

**Figure 3:**
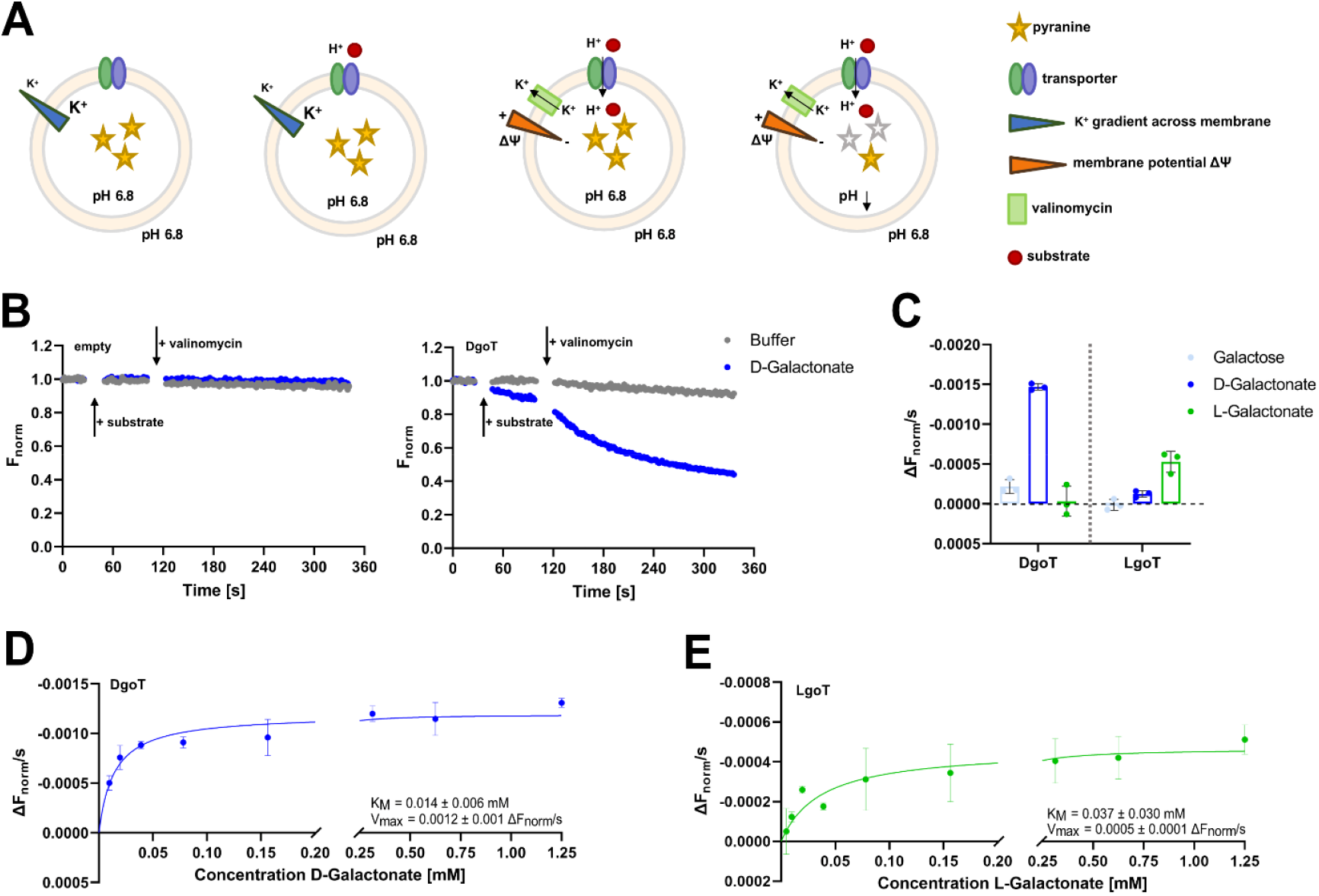
Substrate transport of DgoT and LgoT measured by the pyranine assay. (A) Schematic representation of the pyranine assay. The transporters are reconstituted into liposomes which are filled with the membrane-impermeable fluorescent dye pyranine. While the pH outside and inside the liposome is the same at the beginning, a potassium (K^+^) gradient across the liposome membrane is established, with the higher K^+^ concentration on the inside. The addition of the potassium-selective ionophore valinomycin allows for K^+^ ions to cross the membrane thus, abolishing the K^+^ gradient but simultaneously creating a charge gradient across the membrane (membrane potential ΔΨ). In the presence of a substrate, the proton-coupled transporters utilize this gradient to transport substrate and protons into the liposome. This proton influx lowers the pH inside the liposomes leading to a change in pyranine fluorescence. (B) Transport curve of liposomes without reconstituted protein (empty) and DgoT reconstituted liposomes as an example of a pyranine assay measurement. Blue curves were recorded with addition of the substrate D-galactonate while grey curves represent control recordings with the addition of buffer. The approximate time point of substrate (buffer, D-galactonate) or valinomycin addition is indicated by arrows. Because addition of substrate and valinomycin requires the fluorescence reading to be paused, there are no data points recorded during these timespans. (C) Transport of sugar acids by DgoT and LgoT shown as change in fluorescence over time (ΔF_norm_/s) DgoT shows transport of D-galactonate and LgoT of L-galactonate. Galactose was used as control. Final substrate concentration was 2.5 mM. Error bars correspond to the standard deviation (n = 3). (D-E) Concentration dependent substrate transport of (D) D-galactonate by DgoT and (E) L-galactonate by LgoT with apparent K_M_ and V_max_. Error bars correspond to the standard deviation (n = 2).

Control experiments confirmed the specificity and robustness of the assay. Empty liposomes did not exhibit fluorescence changes under any condition. Likewise, proteoliposomes showed no strong signal change upon addition of substrate only, consistent with the requirement for both substrate and membrane potential to drive proton symport (Figure 3B). Using this system, we evaluated the two *E. coli* ACS homologs, LgoT and DgoT, against the sugar acids identified in the binding assays or assigned by the *in vivo* growth assay. The obtained transport profiles were sharply substrate-specific. DgoT allowed the expected proton-coupled uptake of D-galactonate (K_M_= 14 ± 6 µM), whereas LgoT transported only L-galactonate (K_M_= 37 ± 30 µM), confirming the stereochemical selectivity and consistent with the opposing enantiospecificity previously established for DgoT (Figure 3C-F). Overall, the liposome assays provide a functional readout that resolves ambiguities inherent to ligand binding alone. Across the ACS family, DgoT and LgoT select for the D- and L-galactonate enantiomers, respectively.

### X-ray structure of LgoT reveals an inward facing state

To gain molecular insight into the structure of LgoT and the basis of its stereochemical selectivity, we determined its X-ray structure using lipidic cubic phase crystallization. Crystals diffracted to a resolution of 2.2 Å and belonged to space group C2, with one molecule in the asymmetric unit (ASU) (Table 2). The resulting electron density map was of high quality and enabled model building for most of the protein chain. However, several residues at the N- and C-termini, as well as a short loop region, lacked interpretable density and were therefore not modeled, likely reflecting conformational flexibility.

**Table 2:** Data-collection and refinement statistics.

|  | Native |
| --- | --- |
| PDB ID | 9TXM |
| *Data collection* |  |
| Beam size (FWHM $\mu\text{m}$) | $50.6 \times 50.3$ |
| Flux (ph/s) | $1.48 \times 10^{12}$ |
| Energy (keV) | 12.7 |
| Exposure time per frame (ms) | 30 |
| Number of images | 2000 |
| Space Group | C2 |
| Unit cell constants ($\text{\AA}$) | 122.39, 70.34, 61.38 |
| Unit cell angles ($^\circ$) | 90.0, 107.57, 90.0 |
| Resolution ($\text{\AA}$) | 60.24 – 2.21 (2.24 – 2.21) |
| $R_{\text{merge}}$ | 0.061 (0.401) |
| $R_{\text{meas}}$ | 0.071 (0.463) |
| $R_{\text{pim}}$ | 0.035 (0.227) |
| Total number of observations | 96407 (4686) |
| Unique number of observations | 24877 (1184) |
| Mean $I/\text{sd}(I)$ | 16.3 (2.2) |
| Completeness (%) (ellipsoidal) | 98.89 (95.56) |
| Multiplicity | 3.9 (4.0) |
| $CC_{1/2}$ | 0.998 (0.874) |
| *Refinement* |  |
| Resolution range ($\text{\AA}$) | 60.24 – 2.21 |
| Number of reflections | 24860 |
| R<sub>work</sub> / R<sub>free</sub> | 0.1886 / 0.2282 |
| Test set size (%) | 4.95 |
| Wilson B (Å<sup>2</sup>) | 38.19 |
| Clashscore | 9.03 |
| Number of atoms | 3373 |
| Protein | 3263 |
| Water | 89 |
| Others | 21 |
| *R.m.s. deviations* |  |
| Bonds (Å) | 0.007 |
| Angles (°) | 0.808 |
| *Ramachandran Plot* |  |
| Outliers (%) | 0.24 |
| Allowed (%) | 0.96 |
| Favoured (%) | 98.80 |
| *mean B-factors* |  |
| Overall | 41.21 |
| Protein | 40.92 |
| Water | 47.31 |
| Other | 60.67 |

Values in parentheses are for the highest-resolution shell.

LgoT adopts the canonical major facilitator superfamily (MFS) fold, consisting of twelve transmembrane helices arranged into two pseudo-symmetrical six-helix bundles. Transmembrane helices TM1-TM6 form the N-terminal bundle, whereas TM7-TM12 constitute the C-terminal bundle, with the substrate-binding cavity located at the interface between the two domains. On the cytoplasmic side, the bundles are connected by two intracellular helices (ICH1 and ICH2). A short stretch of two residues linking ICH1 and ICH2 could not be modeled due to insufficient electron density (Figure 4A). The structure captures LgoT in an inward-open conformation, characterized by an open cytoplasmic vestibule and a closed periplasmic side. This conformation closely resembles that of the previously reported wild type DgoT structure (PDB ID: 6E9N) and the AlphaFold2 model of LgoT (AF-P39398-F1-v6). Closure of the periplasmic side is stabilized by a network of salt bridges linking both bundles, typical for the inward-open state in MFS transporters (Figure 4B).

**Figure 4:**
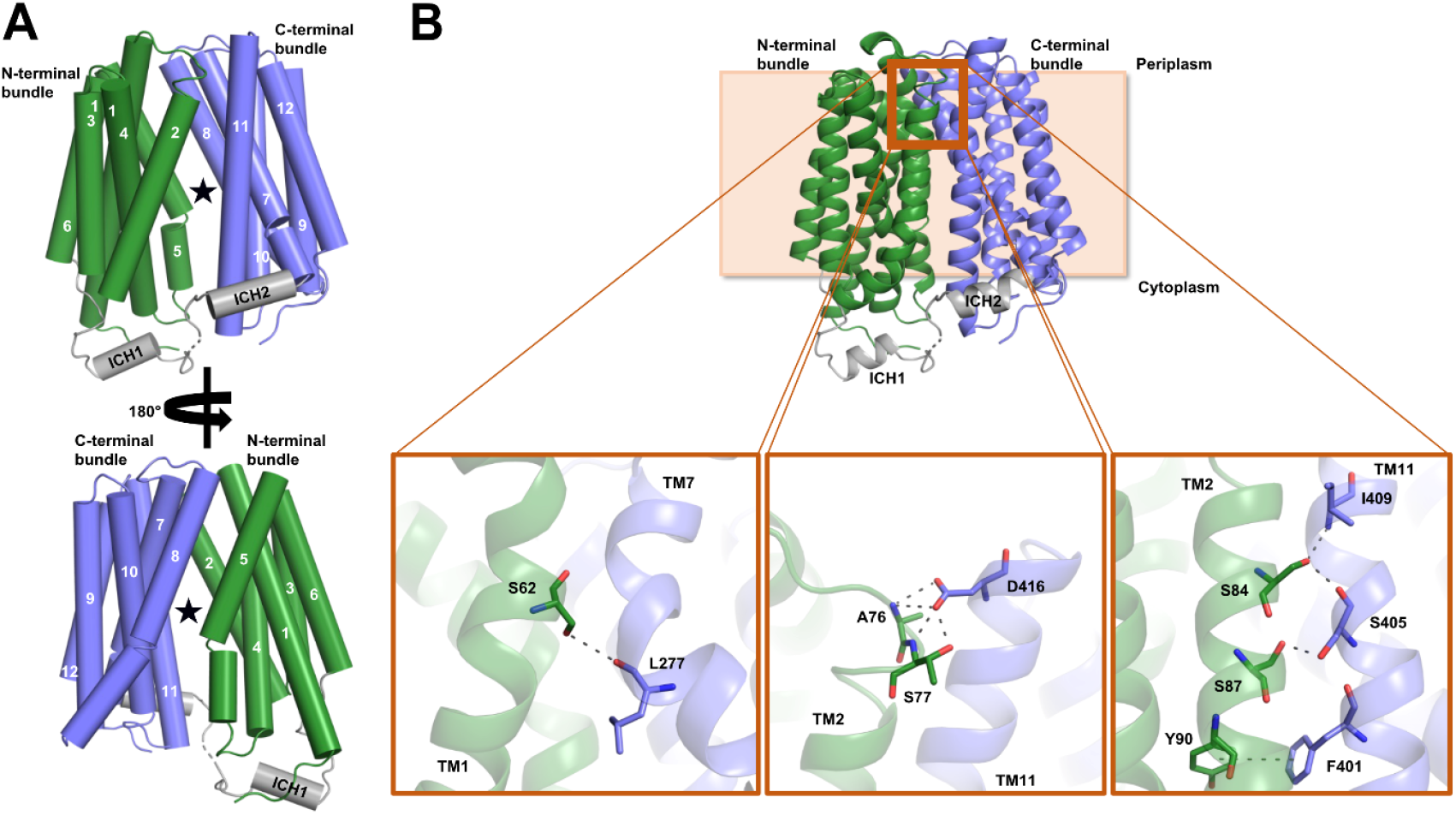
X-ray structure of wild type LgoT from *E. coli*. (A) Atomic model of LgoT in the inward-open state, highlighting the different structural elements of ACS transporters. LgoT adopts the canonical MFS fold consisting of 12 transmembrane α-helices (TMs) which are arranged in two domains of six consecutive helices each, the N-terminal and the C-terminal bundle. The substrate binding site is located between the two bundles. On the cytoplasmic side a linker region connects the N- and C-terminal bundles. For LgoT this linker region includes two intracellular helices (ICH). LgoT is shown as tubes with the N-terminal bundle in green, the C-terminal bundle in blue and the ICH domain in grey. The substrate binding site is marked by a star. TMs are numbered starting from the N-terminus. (B) Interactions between the N- and C-terminal bundle of LgoT in the inward-open state. LgoT is shown as cartoon with the N-terminal bundle in green, the C-terminal bundle in blue and the ICH domain in grey. Insert panels show a close up on labeled residues involved in interactions between the N-terminal and C-terminal bundle on the periplasmic side of the transporter. Potential hydrogen bonds ≤ 3.4 Å and a potential pi-interaction between Y90 and F401 are highlighted by dashes.

Despite extensive efforts, including substrate soaking and co-crystallization with L-galactonate, only the apo structure of LgoT was obtained, consistent with the assumption that the inward-open state represents a substrate-release state.

Because ACS transporters operate as proton-coupled symporters, we examined whether residues implicated in proton coupling are conserved. Indeed, these residues are highly conserved across family members (SI Figure 6), and mutation of the protonation site E144 in LgoT (corresponding to E133 in DgoT) abolishes transport activity (SI Figure 7), supporting a shared coupling mechanism. Structural comparison of LgoT with DgoT and its AlphaFold2 model [55] revealed only minor differences, reflected in low C_α_ r.m.s.d. values of 0.7–2.0 Å over 382 aligned residues (excluding the flexible ICH loop; SI Table 1), indicating a high degree of structural conservation.

### Conserved but Distinct Substrate-Binding Sites Across ACS Transporters

Given the strong global conservation, we next examined whether differences in substrate specificity could be explained at the level of the binding pocket. A comparison of LgoT with DgoT (Figure 5A) shows that most residues previously identified as critical for substrate coordination in DgoT are conserved in LgoT. Consistent with this, electrostatic surface analysis reveals a predominantly positively charged binding cavity in LgoT (Figure 5B), in agreement with its role in transporting a negatively charged sugar acid. Additional non-protein electron density was observed within this cavity; based on its size and shape, it was modeled as glycerol molecule, likely originating from purification or crystallization conditions rather than representing the native substrate.

**Figure 5:**
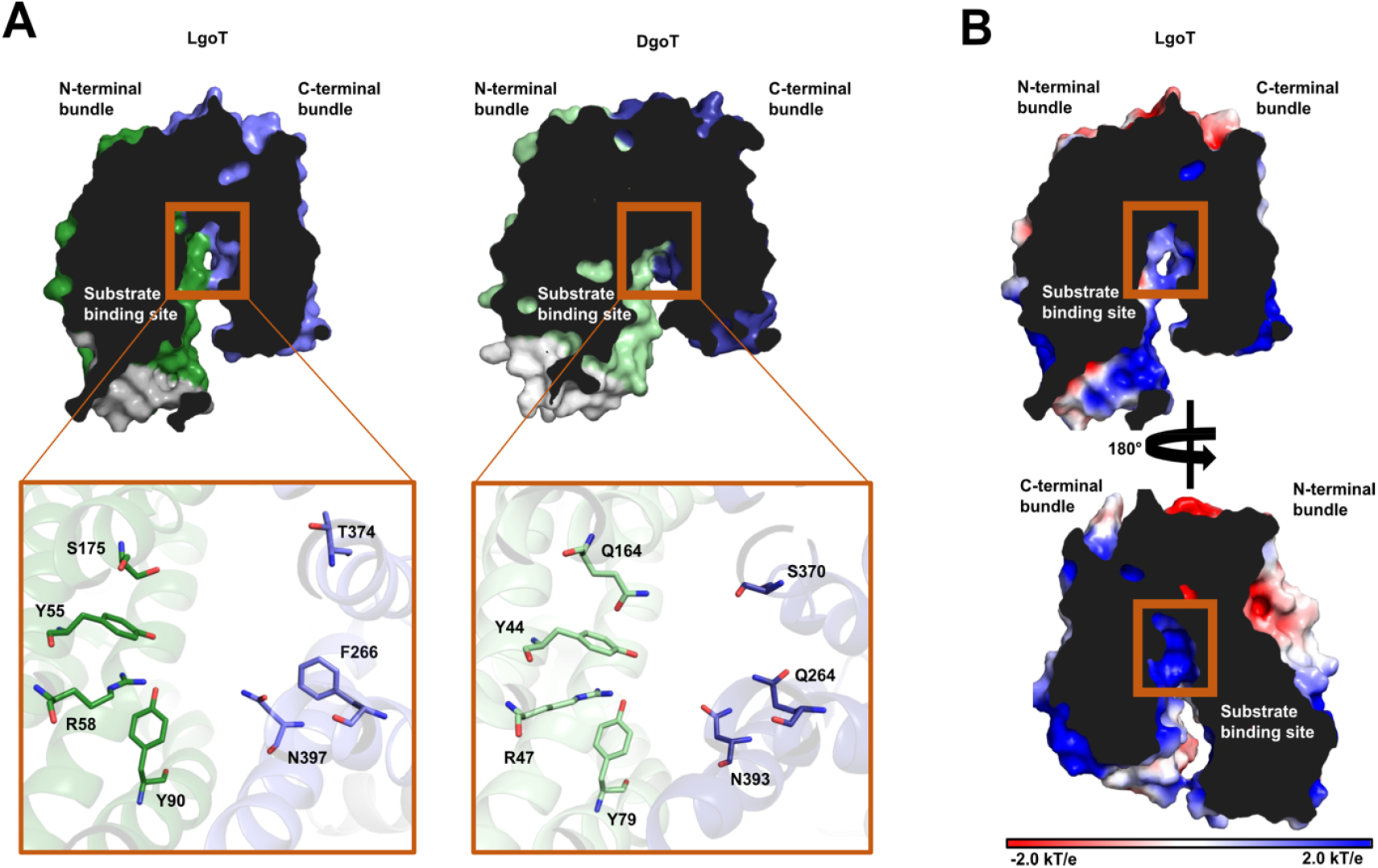
Comparison of the substrate binding site of LgoT and DgoT. (A) Surface representation of LgoT and DgoT. The N-terminal domain is colored in green, the C-terminal domain in blue and the ICH domain in grey. Insert panels show a close up on the substrate binding site, with residues involved in substrate binding in DgoT [21] and the equivalent positions in LgoT shown as stick model. (B) Surface representation of LgoT coloured by electrostatic potential showing the overall positive potential in the binding site. The colour scale is in units of kT/e with blue denoting positive charge and red negative. The electrostatic surface was calculated using the APBS plug-in in PyMOL [56]. 180° rotation for better visibility of the substrate binding site.

To systematically compare substrate coordination across members of the ACS family, we used the DgoT structure in complex with D-galactonate (PDB: 6E9O) as a reference. Seven crucial substrate-coordinating residues were identified in DgoT and mapped onto LgoT, GarP, and GudP using combined sequence and structural alignments (termed position 1-7). Corresponding structures (experimental and AlphaFold2 models) were superimposed (Figure 6 A-B). The position of D-galactonate from the outward-open DgoT structure was used as a structural proxy for the substrate position, assuming a broadly conserved binding mode across the family.

**Figure 6:**
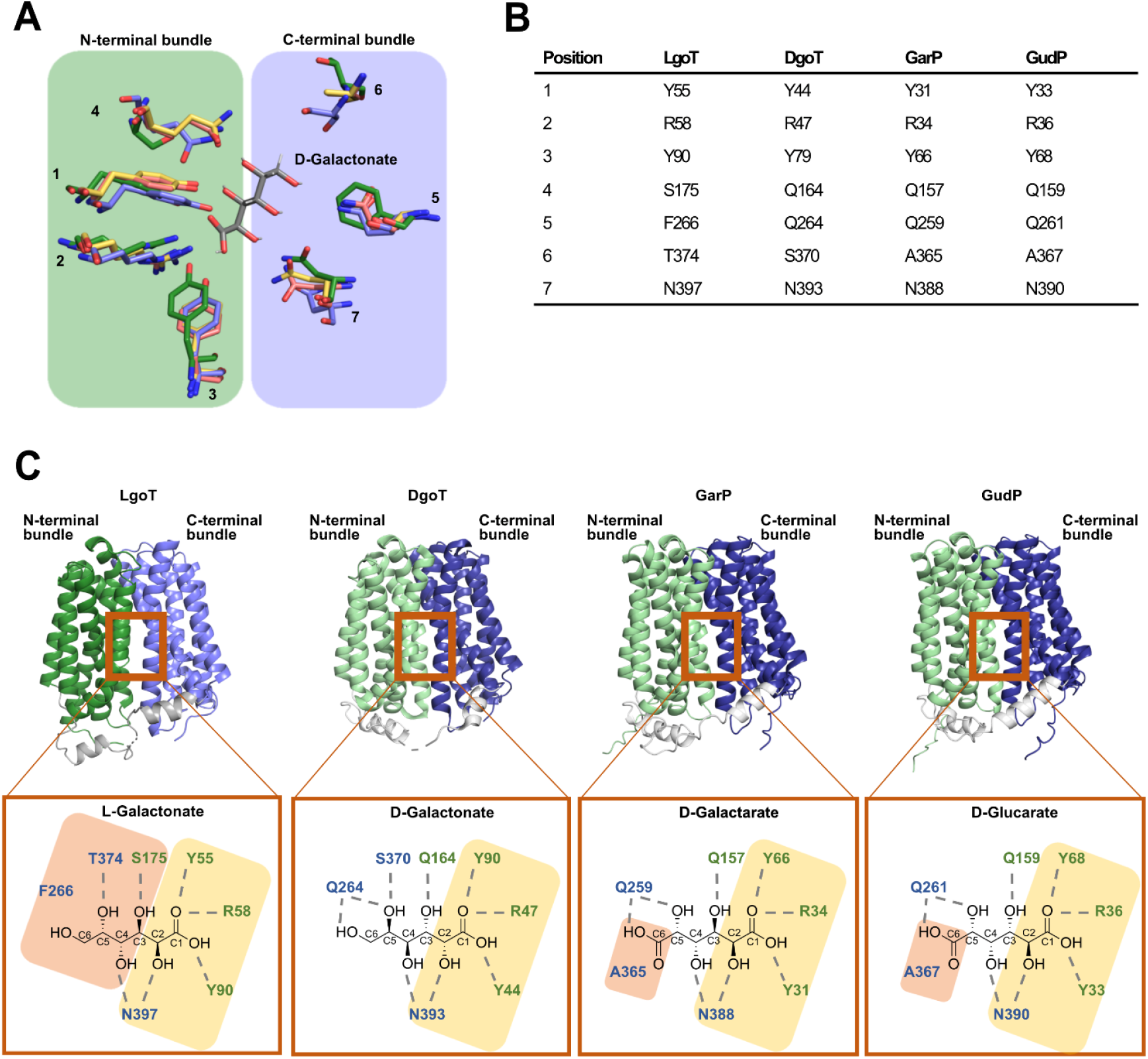
Proposed binding site models of *E. coli* ACS transporters based on multiple sequence alignment. (A) Atomic model of proposed binding site residues from LgoT (this work), DgoT (PDB ID: 6E9N), GarP (AF-P0AA80-F1-v6), GudP (AF-Q46916-F1-v6) in the inward-open state superimposed with the D-galactonate bound to DgoT in the outward-open state (PDB-ID: 6E9O). LgoT residues are shown in green, DgoT in blue, GarP in red and GudP in yellow, D-galactonate is highlighted in gray. Residues of the four transporters belonging to the N-terminal bundle are colored with a green box, residues belonging to the C-terminal bundle are highlighted with a blue box. Coordinating residues are labeled with position 1-7. (B) Amino acid residues proposed to be involved in substrate coordination for each of the four studied *E. coli* ACS transporters. Equivalent residues according to multiple sequence alignment of the protein sequence. Positions analogue to panel A. (C) Atomic models of *E. coli* ACS transporters showing the proposed coordination of substrates in the binding site. LgoT, DgoT (PDB ID: 6E9N), GarP (AF-P0AA80-F1-v6), GudP (AF-Q46916-F1-v6) are all shown in their inward-open state. N-terminal bundles shown in green, C-terminal bundles shown in blue and ICHs domains shown in gray. Residues conserved in all four transporters are highlighted with a yellow box, residues differing between the transporters are highlighted with an orange box.

This analysis reveals a conserved core of interactions across all four transporters. Residues corresponding to positions 1–3 and 7 coordinate the C1 carboxyl group and the C2 hydroxyl group of the substrate and are highly conserved. Given that all four substrates share a C1 carboxylate and a C2 hydroxyl group (albeit with differing stereochemistry), this conservation indicates a shared anchoring mechanism that accommodates both L- and D-sugar acid configurations at C2.

In contrast, key differences are observed at position 6 within the binding site. In DgoT and LgoT, this position is occupied by polar residues (serine and threonine, respectively), that are positioned to interact with the C5 hydroxyl group of D- and L-galactonate. In GarP and GudP, this residue is replaced by alanine, effectively removing side-chain hydrogen bonding capacity and creating additional space within the binding pocket. This change likely accommodates the additional carboxyl group at C6 present in D-galactarate and D-glucarate, substrates of GarP and GudP, but absent in D- and L-galactonate. Thus, variation at this position appears to reflect adaptation to substrate size and charge distribution.

LgoT exhibits the most pronounced differences within the binding site, particularly at positions 4–6. These changes likely underpin its strict preference for L-galactonate, in contrast to the D-specificity of DgoT. While the overall coordination framework is conserved, the mirrored orientation of hydroxyl groups in L-versus D-galactonate would require adjustments in residue identity and positioning to maintain optimal hydrogen bonding geometry. Consequently, changes in side-chain identity and spacing at these positions therefore provide a plausible structural basis for stereoselectivity. The role of residue F266 in LgoT remains speculative in the absence of a substrate-bound structure.

Taken together, the sequence and structural analysis identifies a conserved core of substrate coordination across ACS transporters, combined with transporter-specific adaptations that accommodate differences in substrate stereochemistry and functional groups. While these data provide a coherent model for substrate recognition within the ACS family, definitive mechanistic understanding will require high-resolution structures of substrate-bound states for each transporter.

## Conclusions

This study combines functional assays and structural analysis to define substrate recognition within the *E. coli* ACS transporter family. *In vivo* growth phenotypes together with biochemical and transport measurements establish distinct substrate specificities for all characterized members, including stereoselective transport of L- and D-galactonate by LgoT and DgoT, respectively. The 2.2 Å structure of LgoT reveals an inward-open MFS conformation and, together with comparative modeling across the family, provides a structural framework for understanding conserved features of substrate coordination and the molecular basis of specificity. Collectively, these findings clarify how closely related ACS transporters achieve functional diversification and establish a basis for future mechanistic studies of proton-coupled organic anion transport

## Materials and Methods

### Chemicals

Unless specified otherwise, chemicals were purchased from Sigma-Aldrich, phospholipids were purchased from Avanti Polar Lipids, Inc., n-dodecyl-ß-D-maltoside (DDM), n-decyl-ß-D-maltoside (DM), n-Dodecyl-N,N-Dimethylamine-N-Oxide (LDAO), and n-Nonyl-β-D-Maltopyranoside (NM) from Anatrace, antibiotics and iso-propyl-ß-d-thiogalactopyranoside (IPTG) from Roth, DNase I from Appli-Chem, Lysozyme and protease inhibitor cocktail from Roche, Calcium-D-galactonate hydrate, L-Galactono-1,4-lactone were purchased from Carbosynth, and Terrific broth (TB) from Melford.

### Protein constructs

ACS family members from *E. coli* (Uniprot IDs: ExuT P0AA78, RhmT P76470, DgoT P0AA76, LgoT P39398, GarP P0AA80, GudP Q46916) were cloned into the pNIC-CTHF vector (Addgene plasmid-ID Plasmid #26105) using LIC cloning [57] and the constructs confirmed by DNA sequencing.

### Protein expression and Purification

The transporters were expressed and purified according to established procedures [58, 59]. In short, after transformation into *E. coli* strain C41 (DE3) cells were grown in TB medium supplemented with 30 μg/mL kanamycin at 37 °C. At an OD_600nm_ of 0.6 the cells were induced by 0.2 mM IPTG incubated at 18°C for a further 16 hours, and harvested by centrifugation (9379 × g, 15 min, 4 °C). The cells were resuspended in lysis buffer (20 mM sodium phosphate pH 7.5, 300 mM NaCl, 5% (v/v) glycerol, 15 mM imidazole, 0.5 mM TCEP, 5 units/mL of DNase I, 1 mg/mL lysozyme, and protease inhibitor) at a ratio of 5 mL lysis buffer per 1 g of cells. Cell lysis was performed by three cycles using an EmulsiFlex-C3 (Aventin) at 10,000 psi and undisrupted cells and debris removed by a low-speed centrifugation step (10,000 × g, 15 min, 4 °C). The cell membranes were pelleted by ultracentrifugation (142,400 × g, 50 min, 4 °C), resuspended in lysis buffer, supplemented with 1% DDM, 0.5 mM TCEP and protease inhibitor, and stirred for 1 h at 4 °C. For purification using Immobilized Metal Affinity Chromatography (IMAC), the supernatant of an additional ultracentrifugation step (104,600 × g, 50 min, 4 °C) was applied to Ni-NTA agarose resin (ThermoFisher) and incubated for 1h at 4 °C on a rotating wheel. After washing with increasing imidazole concentration (20 mM HEPES pH 7.0, 100 mM NaCl, 5% (v/v) glycerol, 15-25 mM imidazole, 0.03% (w/v) DDM, 0.5 mM TCEP) the protein was eluted with elution buffer (20 mM HEPES pH 7.0, 100 mM NaCl, 5% (v/v) glycerol, 200 mM imidazole, 0.03% (w/v) DDM, 0.5 mM TCEP), supplemented with TEV protease, and dialysed overnight at 4 °C against SEC buffer (20 mM HEPES pH 7.0, 100 mM NaCl, 5% (v/v) glycerol, 0.03% (w/v) DDM, 0.5 mM TCEP). The cleaved protein was recovered via negative IMAC, concentrated to 5 mL using a 100 MWCO concentrator (Corning Spin-X UF concentrators) and applied to a HiLoad® 16/600 Superdex® 200 pg (GE Healthcare Life Sciences) on an ÄKTA Pure system (GE Healthcare Life Sciences) equilibrated in SEC buffer with either 0.03 % (w/v) DDM, 0.3 % (w/v) DM, 0.4 % (w/v) NM, or a mixture from 0.03 % (w/v) DDM and 0.02 % (w/v) LDAO). Protein-containing fractions were pooled and concentrated to 40 mg/mL for crystallization or 5 mg/mL for functional assays.

### Reconstitution of LgoT into Salipros

LgoT was reconstituted into Salipros as previously described, using a 1:20:35 molar ratio of membrane protein to SapA to lipid [51, 52]. POPS (Avanti Polar Lipids) was used as lipid and incubated at 37 ºC for 10 min, LgoT was added and incubated at RT for 15 min. After addition of SapA, the mixture was incubated at RT for another 20 min. 50 mg of Biobeads (Biorad) were added per 100 μL of sample and the sample was incubated overnight at 4ºC with gentle agitation. The reconstituted membrane protein was recovered by SEC on a Superdex® 200 Increase 10/300 GL column (GE Healthcare Life Sciences) equilibrated with SEC buffer without detergent.

### *In vivo* growth assays

For *in vivo* growth assays, LB medium, for the knockout mutants supplemented with 30 μg/mL kanamycin, was inoculated with either the parental (BW25113) or one of six ACS transporter knockout strains (ExuT, RhmT, DgoT, LgoT, GarP and GudP) and grown at 37 °C overnight. These pre-cultures were used to inoculate M9 minimal medium supplemented with 2 g/L glucose, and 30 μg/mL kanamycin for the knockout mutants, to an OD_600nm_ of 0.05. After incubation at 37 °C overnight, cells were harvested by centrifugation (3163 × g, 5 min, 4 °C), washed twice with 1 × PBS, and resuspended to OD_600nm_ of 10 in 1 × PBS. The cell suspension was spotted on Minimal agar plates supplemented with 2 g/L per carbon source and incubated at 37 °C overnight.

### Ligand binding assay

To monitor ligand binding of ACS transporters by nanoDSF, proteins were diluted to 0.5 mg/mL in nanoDSF buffer (200 mM HEPES pH 7.0, 100 mM NaCl, 5% (v/v) glycerol, 0.03% (w/v) DDM) and incubated with ligands for 10 min at RT at a final ligand concentration of 2.5 mM. For ligand binding with different detergents, the proteins were diluted to 0.5 mg/mL in nanoDSF buffer with either 0.03 % (w/v) DDM, 0.3 % (w/v) DM, 0.4 % (w/v) NM, or a mixture from 0.03 % (w/v) DDM and 0.02 % (w/v) LDAO) or no detergent for POPS Salipros. For ligand binding at different pH, the proteins were diluted to 0.5 mg/mL in sodium phosphate nanoDSF buffer (200 mM sodium phosphate pH 6.0-8.0, 100 mM NaCl, 5% (v/v) glycerol, 0.03% (w/v) DDM). Samples were loaded into standard grade nanoDSF capillaries (Nanotemper) and thermal denaturation measured with a Prometheus NT.48 device (Nanotemper) controlled by PR. ThermControl (version 2.1.2). Excitation power was adjusted to 10% and samples were heated from 20 °C to 90 °C with a slope of 1 °C/min. The results were analysed with Excel (Microsoft) and GraphPad Prism 9.5.1 (GraphPad Software, CA, USA).

### Liposome reconstitution

To prepare liposomes, POPE and POPG (Avanti Polar Lipids) were mixed in a 3:1 (w/w) ratio, dissolved in chloroform, washed twice with pentane and resuspended into degassed liposome buffer (50 mM potassium phosphate pH 7.0) to a final lipid concentration of 20 mg/mL. After three freeze-thaw cycles liposomes were diluted to 5 mg/mL with liposome buffer and extruded 11 times through a LiposoFast Liposome factory 400 nm filter (Avestin, Inc.). For reconstitution, proteins were purified as described above but with 0.3% DM instead of 0.03% DDM and used without further concentration after SEC. Fractions containing at least 0.5 mg/mL protein were mixed with extruded liposomes in a 1:60 or 1:30 protein-to-lipids ratio and incubated for 1 h at RT. Samples were dialysed for 2 h at 4°C against excess of liposome buffer using SnakeSkin Dialysis Tubing (ThermoFisher) with a MWCO of 10 kDa. After three buffer exchanges over the next 36 h, liposomes were harvested by centrifugation (100,000 × g, 30 min, 4 °C). Empty liposomes as control samples were prepared by substituting the protein solution with SEC buffer with 0.3% DM. Liposomes were subjected to two freeze-thaw cycles and stored at -80 °C until further use. The reconstitution efficiency was determined using SDS-PAGE densitometry, comparing known concentrations of the protein to the corresponding proteoliposomes.

### Transport assay

Proteoliposomes were thawed at RT and after centrifugation (100,000 × g, 30 min, 4 °C) the pellet was resuspended in pyranine inside buffer (5 mM HEPES pH 6.8, 120 mM KCl, 2 mM MgSO_4_) supplemented with 10 mM pyranine. To form unilamellar vesicles, 7 freeze-thaw cycles were performed and the liposomes 11 times extruded through a LiposoFast Liposome factory 400 nm filter (Avestin, Inc.). After centrifugation (60,000 × g, 30 min, 15 °C) the pellet surface was gently washed with pyranine inside buffer and the pellet subsequently resuspended into pyranine inside buffer. After removing excess pyranine dye using a G-25 spin column, the liposomes were harvested by centrifugation (60,000 × g, 30 min, 15 °C), and resuspended into pyranine inside buffer to a final protein concentration of 1.25 mg/mL. For the uptake assay, 4 μL liposome solution was diluted in 200 μL pyranine outside buffer (5 mM HEPES pH 6.8, 120 mM NaCl, 2 mM MgSO_4_), establishing a potassium gradient across the liposome membrane. After 30 s, substrate was added to a final substrate concentration of 2.5 mM and quickly mixed before the measurement was continued, water was used for control. All substrate stocks were prepared in water to reach a final concentration of 50 mg/ml. To dissipate the potassium gradient and start the transport reaction by creating a proton-gradient with negative charge inside the liposomes, valinomycin was added after 50 s to a final concentration of 1 μM. The transport activity is monitored via the pyranine fluorescence exciting at a wavelength of 415 nm and 460 nm and measuring emission at a wavelength of 510 nm with a Tecan Spark® 20 M Microplate reader (Tecan Trading AG) using a 96-well black chimney plate (Greiner). As controls, empty liposomes were measured on the same day as the protein samples. The results were analysed with Excel (Microsoft) and GraphPad Prism 9.5.1 (GraphPad Software, CA, USA). Fluorescent values at 460 nm were divided by the fluorescent values at 415 nm, and the result again divided by the average fluorescence ratio of the first 28 s before substrate addition to normalize all data. Thus, the relative fluorescence for each sample starts around a value of 1 and is no longer dependent on the total amount of pyranine loaded into the liposomes as this might differ between preparations. The transport signal was determined starting from the time point after addition of valinomycin to the sample. The slope of the sample signal between 150 – 250 s was calculated by linear curve fitting. To correct for unspecific interactions, the slope of the empty liposomes signal was subtracted from the proteoliposome signal. The corrected curve indicates transport of the substrate by the respective protein.

### Crystallization

Two Hamilton syringes connected by an LCP syringe coupler (TTP Labtech) were used to mix LgoT at 40 mg/mL and the lipid Monoolein in a 2:3 (v/v) ratio until the resulting mesophase was clear and stable over time. Crystallisation drops of 80 nL mesophase and 800 nL reservoir buffer (0.1 M Tris-HCl pH 7.0, 0.2 M Li_2_SO_4_, 30% PEG400 (w/v)) were dispensed into a 96-well glass sandwich plate by a Mosquito-LCP robot (TTP Labtech). Crystals grew after 7 days of incubation at 19 °C and were harvested after 14 days.

### X-ray data collection and processing

X-ray data were collected at 100 K on beamline P14 (EMBL-Hamburg, Germany) with an Eiger 16 M detector (Dectris). Data were processed using Xia2 with DIALS [60, 61], Aimless [62], and Pointless [63, 64]. Phaser [65] was used to solve the phase problem by molecular replacement using a previously determined DgoT structure as search model (PDB-ID: 6E9N). Structure refinement was performed using iterative rounds of *phenix*.*refine* [66] and manual model building with COOT [67]. PyMol (Schrödinger LLC) was used to generate figures. Table 2 shows statistics for data collection and refinement. Atomic coordinates and the structure factors for the LgoT structure were deposited in the protein data bank with the following accession number: 9TXM

## Acknowledgements

We thank the Sample Preparation and Characterization facility of EMBL Hamburg for support in this project and the beamlines P13 and P14 at EMBL Hamburg for regular access. We acknowledge the group of Janosch Hennig for NMR analysis of D- and L-galactonate. All past group members are acknowledged for their input to this manuscript. This research was supported through a grant from the German-Israeli Foundation (GIF grant #G-1288-207.9/2015) to CL.

## Declaration of Generative AI and AI-Assisted Technologies

During the preparation of this work, the authors used Grammarly and ChatGPT to improve the style and grammar of the text. After using this tool or service, the authors reviewed and edited the content and take full responsibility for the content of the publication.

