## Supplementary File for "Structural basis of substrate recognition and transport in bacterial ACS transporters"

Christian Löw

European Molecular Biology Laboratory Hamburg, Notkestrasse 85, D-22607 Hamburg, Germany.

Twitter handle: @AllUNeedIsLoew

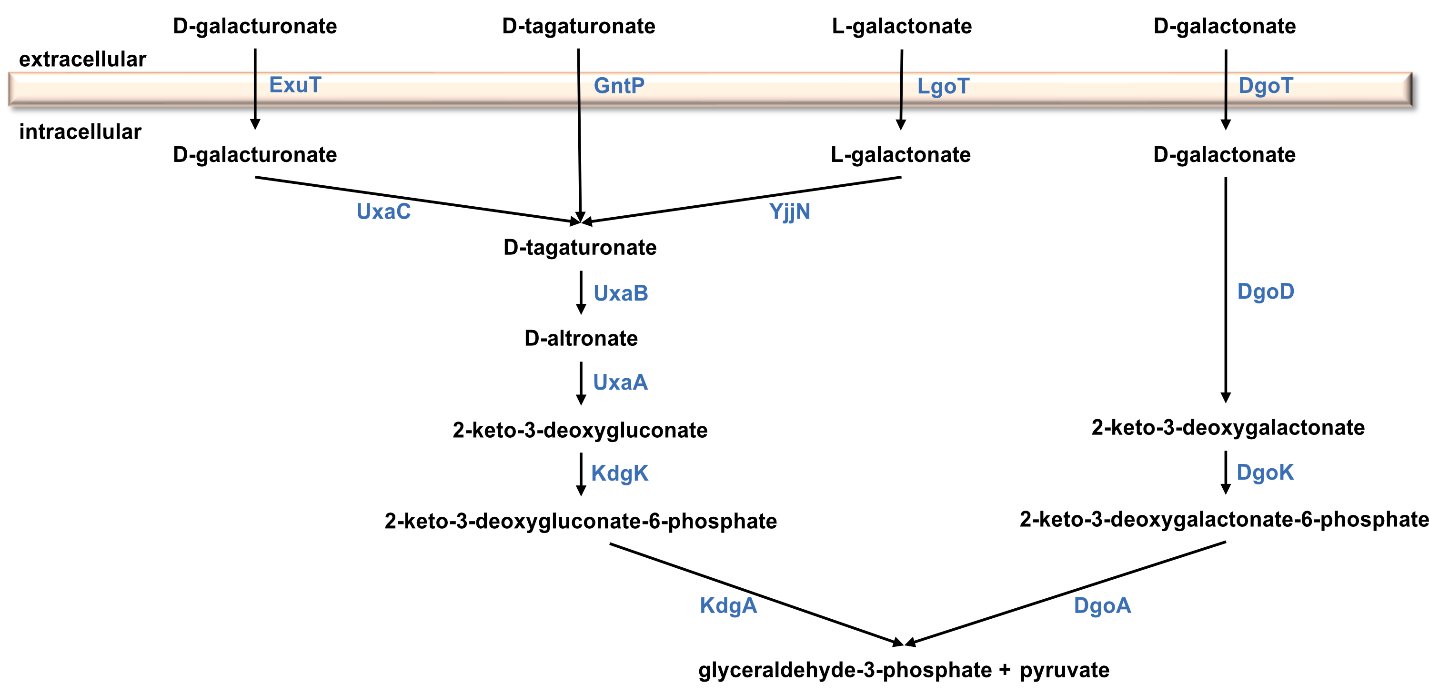

SI Figure 1: Import and utilization of the sugar acids D-galacturonate, D-tagaturonate, L-galactonate and D-galactonate in E. coli via different ACS transporters. Enzymes and breakdown products from different pathways are stated. Figure adapted from [1,2].

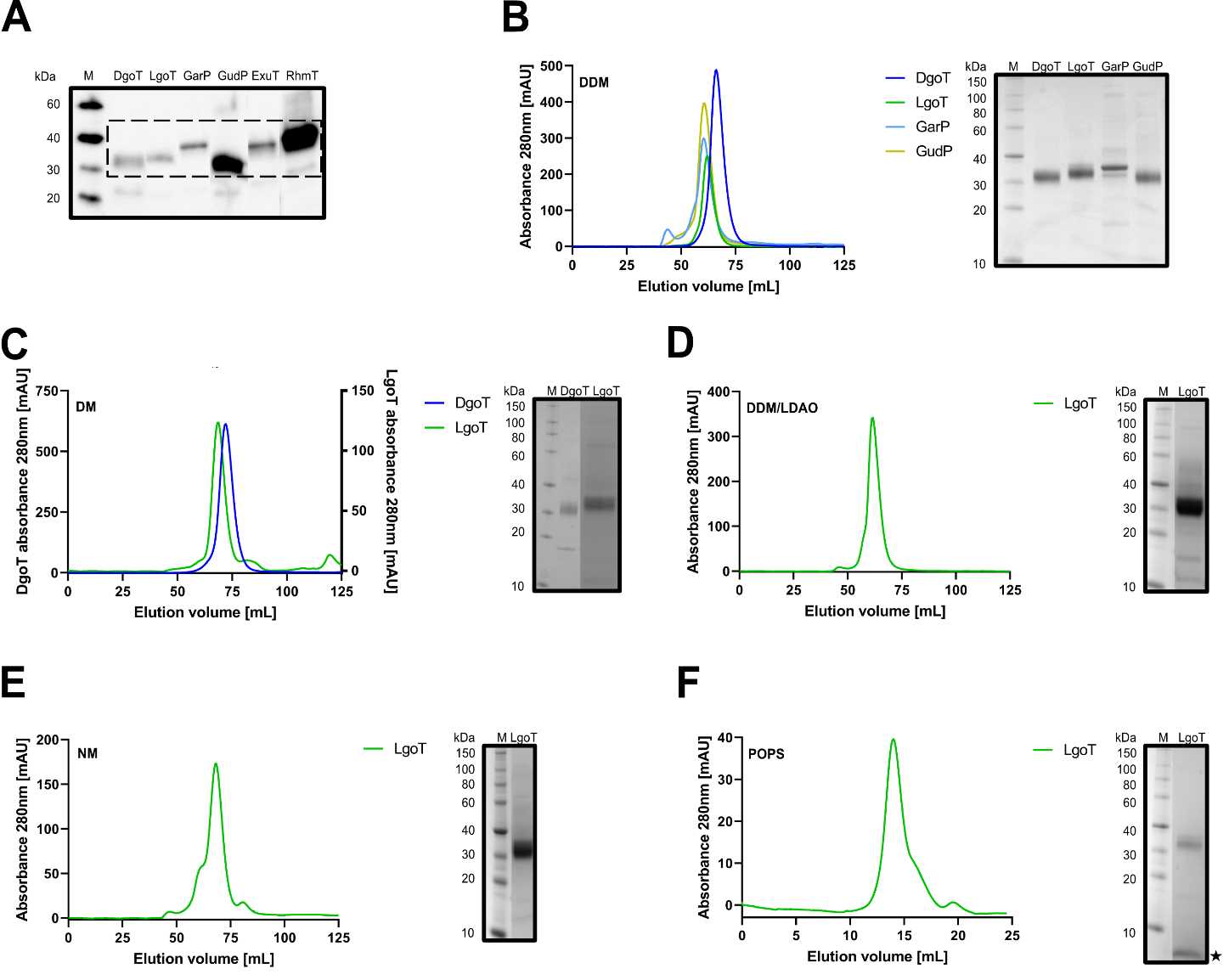

SI Figure 2: Expression and purification of ACS transporters. (A) All six transporters are expressed with an N-terminal His_6_-tag. Cells overexpressing one of the six E. coli ACS transporters were lysed and the soluble lysate was analyzed for the transporter’s presence by immunoblotting using His_6_-specific antibodies. (B-F) Size exclusion chromatography (SEC) profiles of (B) DgoT, LgoT, GarP and GudP in DDM, (C) DgoT and LgoT in DM, (D) LgoT in DDM/LDAO, (E) LgoT in NM and (F) LgoT reconstituted into Salipros with POPS. The star denoted SapA from the Salipros. A HiLoad® 16/600 Superdex® 200 pg column (GE Healthcare Life Sciences) was used in (B-E) and a Superdex® 200 Increase 10/300 GL column (GE Healthcare Life Sciences) in (F). Insert panels show SDS-PAGE gels of the peak fraction. SEC conditions are described in Material and Methods. DgoT is shown in dark blue, LgoT in green, GarP in light blue and GudP in dark yellow.

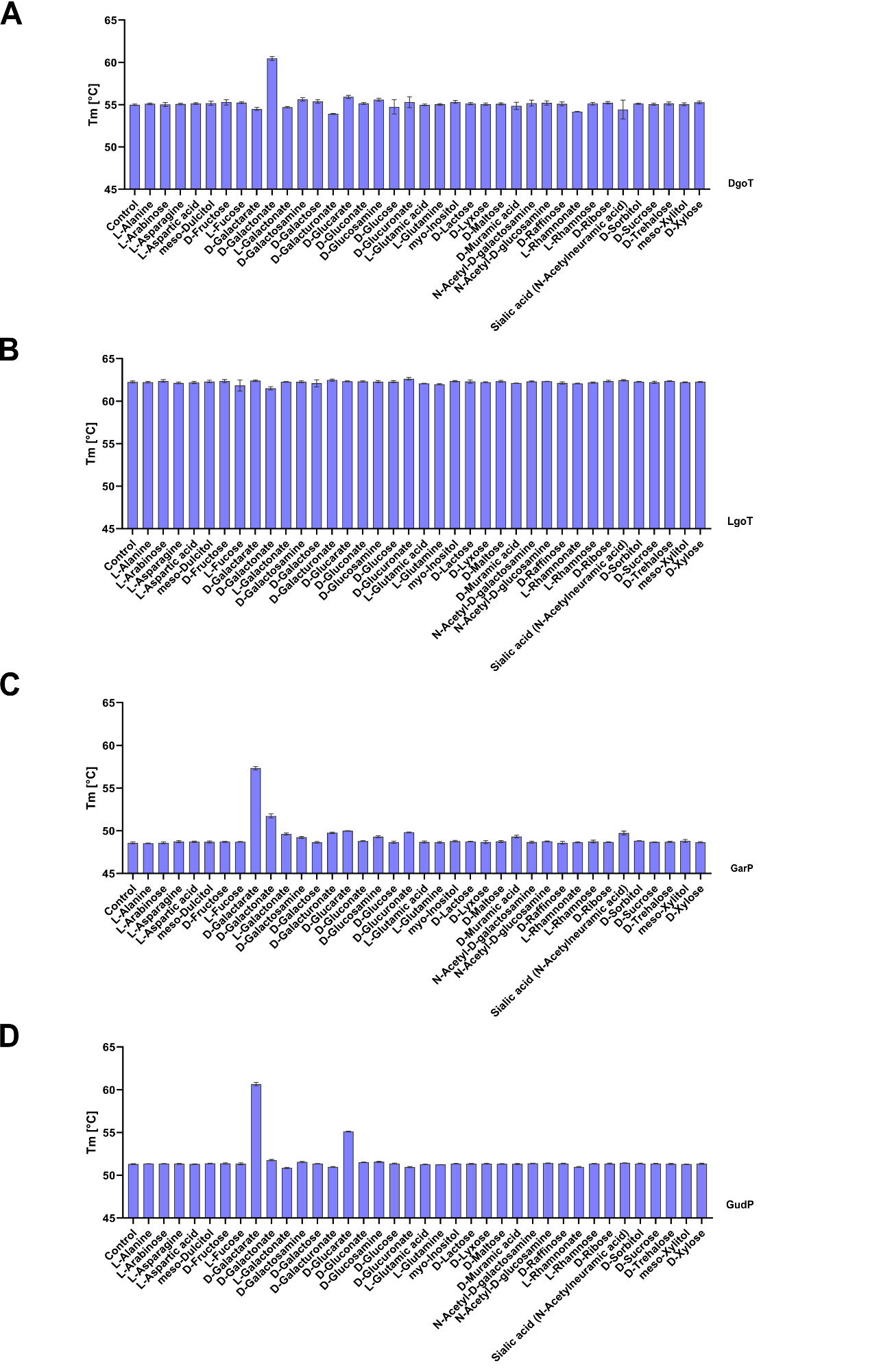

SI Figure 3: Ligand binding screen for ACS transporters measured by nanoDSF. Melting temperature (T_m_) of (A) DgoT, (B) LgoT, (C) GarP and (D) GudP without ligand (control) and upon ligand addition. Protein stability was measured at a protein concentration of 0.5 mg/mL with ligands added to a final concentration of 2.5 mM. The samples were heated from 20 °C to 90 °C. Error bars correspond to the standard deviation (n = 3). Further nanoDSF conditions are described in Material and Methods.

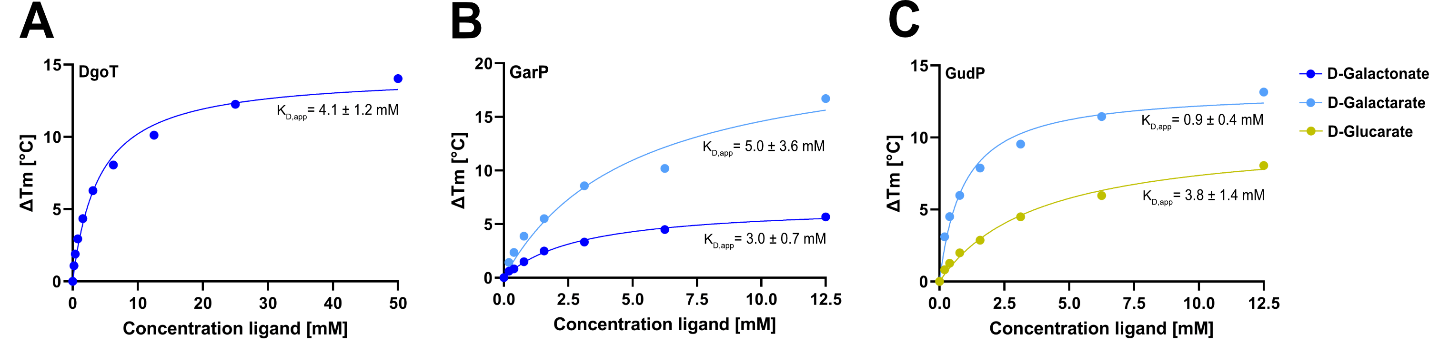

SI Figure 4: Concentration dependent ligand binding of ACS transporters. Difference in T_m_ upon addition of (A) D-galactonate to DgoT, (B) D-galactonate and D-galactarate to GarP and (C) D-galactarate and D-glucarate to GudP with apparent K_D_,_app_. Proteins were measured at a concentration of 0.5 mg/mL. The samples were heated from 20 °C to 90 °C. Further nanoDSF conditions are described in Material and Methods.

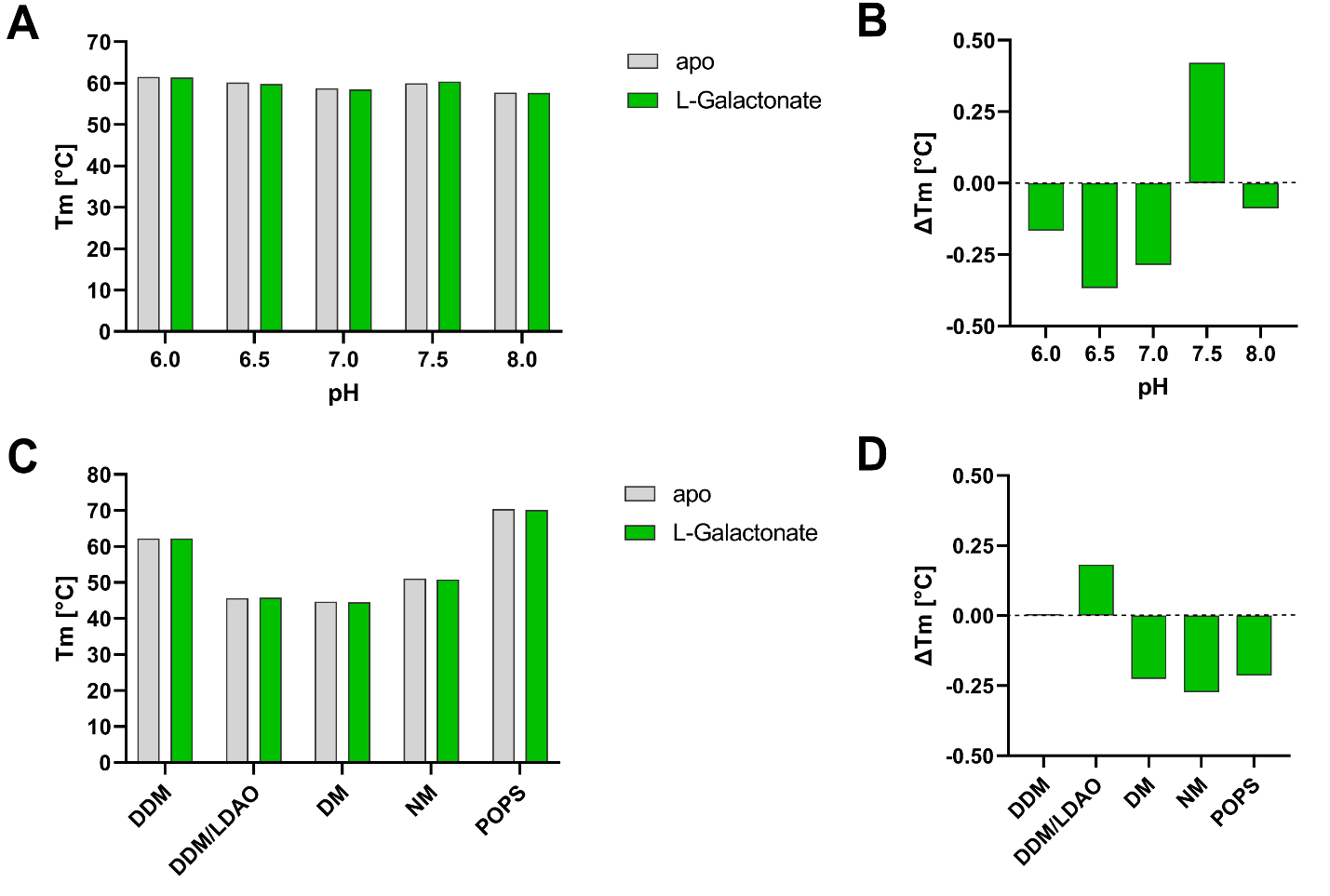

SI Figure 5: Influence of buffer pH and detergent on ligand binding of LgoT measured by nanoDSF. (A) Melting temperature (T_m_) of LgoT in sodium phosphate buffer of different pH without (grey) and with addition of L-galactonate (green). (B) Difference in T_m_ upon addition of L-galactonate to LgoT in sodium phosphate buffer of different pH. (C) Melting temperature (T_m_) of LgoT in solubilized in HEPES buffer pH 7.0 with the different detergents 0.03% (w/v) DDM, 0.03% (w/v) DDM and 0.02% (w/v) LDAO, 0.3 % (w/v) DM, 0.4% (w/v) NM or, Salipros with the lipid POPS without (grey) and with addition of L-galactonate (green). (D) Difference in T_m_ upon addition of L-galactonate to LgoT in the different detergents 0.03% (w/v) DDM, 0.03% (w/v) DDM and 0.02% (w/v) LDAO, 0.3 % (w/v) DM, 0.4% (w/v) NM or, Salipros with the lipid POPS. LgoT was measured at a concentration of 0.5 mg/mL with L-galactonate added to a final concentration of 2.5 mM. The samples were heated from 20 °C to 90 °C. Further nanoDSF conditions are described in Material and Methods.

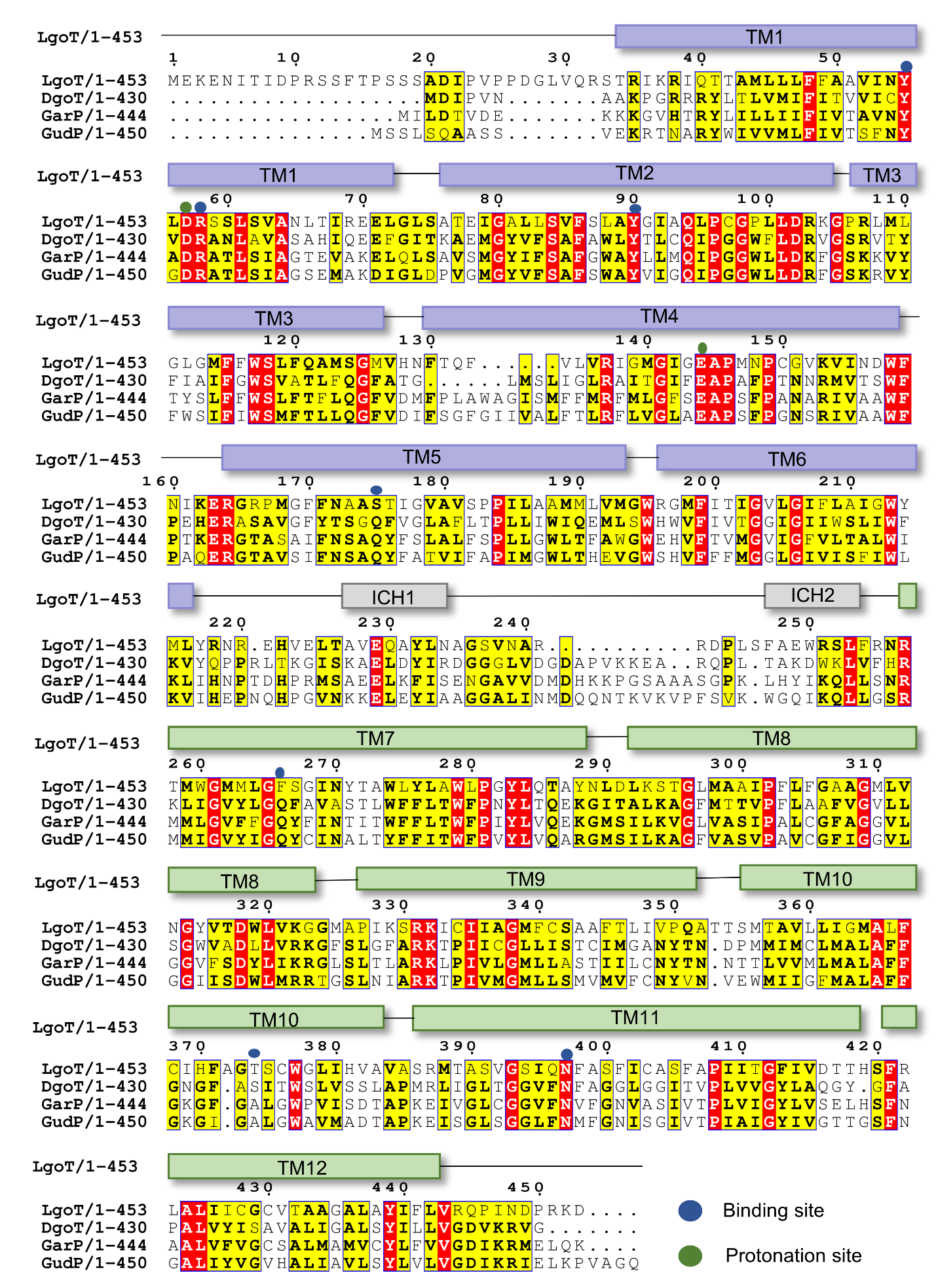

SI Figure 6: Multiple sequence alignment of ACS transporters DgoT, LgoT, GarP and GudP from E. coli. Sequence similarities are colored in yellow for similar and red for identical residues. Numbering is done according to the LgoT sequence. Residues proposed to be important for ligand binding or protonation are marked with blue and green dots respectively. For the LgoT sequence the regions of the 12 TMs of the canonical MFS fold are shown in slate for the N-terminal bundle and green for the C-terminal bundle. The ICH domains position in the LgoT sequence are shown in gray. Both residues predicted to be protonated D57 and E144 are identical in all four proteins as are four out of the seven proposed ligand binding residues, Y55, R58, Y90 and N397. Two more are identical for all but LgoT, namely S175 and F266 which are glutamine residues in the other transporters. Sequence alignment was performed with Jalview and Clustal W [3,4].

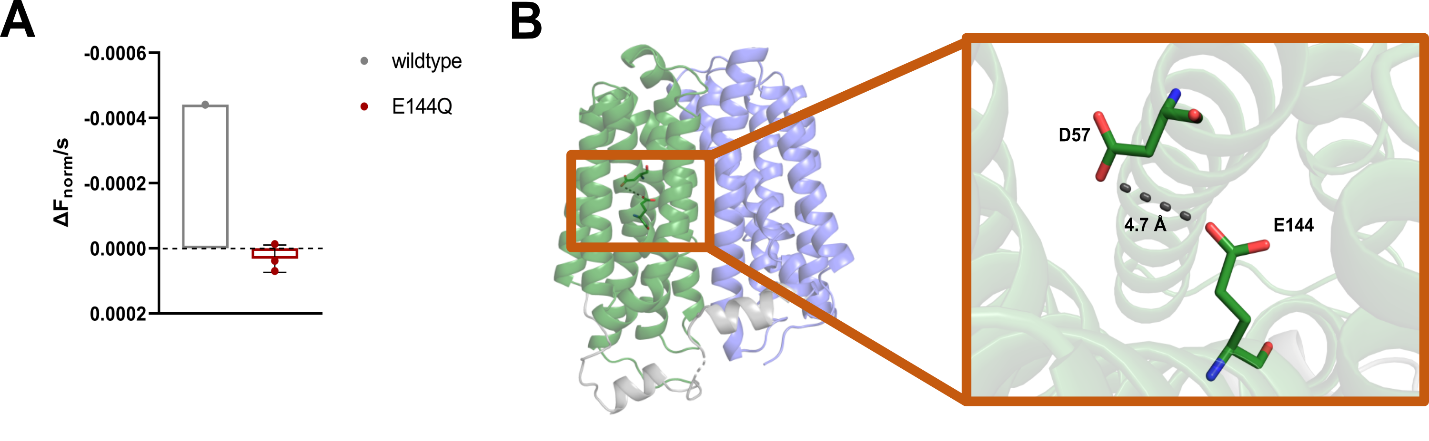

SI Figure 7: LgoT protonation site mutant E144Q. (A) Transport of sugar acids by wildtype and E144Q LgoT shown as change in fluorescence over time (∆F_norm_/s). DgoT shows transport of L-galactonate by wildtype LgoT while the transport is abolished in the E144Q mutant. Final substrate concentration was 2.5 mM. Error bars correspond to the standard deviation (n = 3). (B) Positions of the proposed protonation site residues D56 and E144 in the inward-open state of wildtype LgoT. LgoT is shown as cartoon with the N-terminal bundle in green, the C-terminal bundle in blue and the ICH domain in grey. Insert panels show a close up on labeled residues proposed to be involved in protonation.

SI Table 1: Structure comparison of LgoT, DgoT and the AlphaFold model of LgoT. Cα root mean square deviation (r.m.s.d.) values were calculated over the whole sequence, excluding the ICH loop (L216-N256 for LgoT and V205-H254 for DgoT).

| **PBD-ID** | **Protein** | **Space group** | **Molecules per ASU** | Cα **r.m.s.d. (Å)** | | | |
| --- | --- | --- | --- | --- | --- | --- | --- |
|  |  |  |  | **Intramolecular** | **DgoT (6E9N)** | | **AlphaFold LgoT** |
|  |  |  |  |  | **Chain A** | **Chain B** | **1-453** |
| 9TXM | LgoT | C2 | 1 | n.a. | 2.03 | 1.93 | 0.72 |
| 6E9N | DgoT | P2_1_ | 2 | 0.61 | n.a. | n.a. | n.a. |
